# Skuas mortalities linked to positives HPAIV A/H5 beyond Polar Antarctic Circle

**DOI:** 10.1101/2025.03.02.640960

**Authors:** Fabiola León, Claudia Ulloa-Contreras, Eduardo J. Pizarro, Pablo N. Castillo-Torres, Karla B. Díaz-Morales, Ana Cláudia Franco, Francine C. B. Timm, Miguel L. Corrêa, Lucas Krüger, Elie Poulin, Catalina Pardo-Roa, Juliana A. Vianna

## Abstract

The ongoing extinction crisis, driven by human activity, poses a significant threat to seabirds and it’s especially relevant in highly valuable environments such as Antarctica. Among these threats, seabirds face the risk of local extinctions due to emerging infectious diseases like the Highly Pathogenic Avian Influenza Virus (HPAIV).Progressive spread of HPAIV A/H5N1 outbreaks across South America and the sub-Antarctic islands have been detected, reaching the northern regions of the West Antarctic Peninsula (WAP) during the 2023-2024 season. Here we conducted a comprehensive epidemiological survey conducted on sixteen seabird nesting localities along the WAP from November 2024 to January 2025 to assess the health status of the Antarctic seabirds and detect the presence of HPAIV. We observed unusual mortalities among nesting populations of skuas, with a total of 35 deaths skuas recorded along the WAP and beyond the Antarctic Polar Circle, including Important bird breeding areas around Margarita Bay. HPAIV A/H5 was confirmed in all dead skuas sampled (n=11), from six different locations. This finding represents the southernmost record of seabird mortality in Antarctica related to HPAIV to date. The expansion of HPAIV observed here raises concerns about further spread of avian flu out the Antarctic Peninsula, potentially leading to increased mortality rates in the Antarctic bird populations. These findings are relevant for the assessment of the general health status of Antarctic seabird populations and provide a baseline for the continuous monitoring of the HPAIV spread in avian species during the next breeding seasons.

## Introduction

Seabirds are among the groups of vertebrates with a larger number of species under risk of extinction (Butchart et al., 2004; Días et al., 2019; Croxall et al., 2012; Kruger 2022). While climate change and anthropogenic pressures are on the top of the rank for factors driving extinction risk of seabirds, emerging infectious diseases (EIDs) with “spill-over” are also of significant concern for birds populations, especially for species that breed in densely aggregated colonies (Días et al., 2019; Garces & Pires, 2024; McCallum et al., 2024, Daszac et. al 2000). For instance, infectious diseases such as avian cholera have significantly affected the demography of colonies of Indian yellow-nosed albatross (*Thalassarche carteri*) and *Diomedea* albatrosses (Rolland et al 2009, Jaeger et al., 2018, 2020, Ventura et al., 2021).

Currently, Highly Pathogenic Avian Influenza Virus (HPAIV) is one of the most concerning pathogens, already significantly impacts on seabird species (FAO, 2023; BirdLife International, 2024; Garcés & Pires, 2024). In 2021-2022 an outbreak of H5N1 HPAIV was detected in the North Hemisphere (Wille and Barr, 2022) and expanded globally within a few months, causing mortality events in wild birds, (Wille and Barr, 2022; Ramey et al., 2022; Puryear et al., 2024). Between November and December 2022, the first cases of HPAIV in South America birds was reported (Fernández-Díaz et al., 2023; Ruiz-Saenz et al., 2023; Ministerio Salud Perú 2022, 2023). Its arrival in South America was unprecedented, affecting more than 14 orders of wild birds, several of which are classified as endangered, vulnerable or near-threatened (Gamarra-Toledo et al., 2023; Azat et al., 2024; Godoy et al., 2023; Pardo-Roa et al., 2023). Over 20,000 wild bird deaths were reported in Perú due to HPAIV within just four weeks, with Peruvian pelicans (*Pelecanus thagus*) and Peruvian boobies (*Sula variegata*) being the most affected species (Fernández-Díaz et al., 2023; Ministerio Salud Perú 2022, 2023). In Chile, the same two species were also among the affected. The orders that present the higher mortality rates were Suliformes (Peruvian Booby and Guanay Cormorant), Pelecaniformes (Peruvian pelicans,Pelicanus thagus), Anseriformes (Black-necked Swan, *Cygnus melancoryphus*) and Charadriiformes (*Larus* sp.) (Pardo-Roa et al., 2023; Ariyama et al., 2023). In Peru, Chile and Argentina it caused mass mortality events of *Otaria flavescens* as well (Uhart et al., 2024; Ulloa et al., 2023). Unexpected mortality events of seabirds in South Georgia and the Falkland/Malvinas Islands, with signs consistent with HPAIV, were subsequently confirmed through RT-qPCR, and identified as HPAIV A/H5N1 subtype 2.3.4.4b (Banyard et al., 2024). At that time, Brown Skuas (*Stercorarius antarcticus*), Kelp Gulls (*Larus dominicanus*), Gentoo Penguins *(Pygoscelis papua*), Southern Rockhopper Penguins (*Eudyptes chrysocome*), Black-browed Albatrosses (*Thalassarche melanophris*), Wandering Albatrosses (*Diomedea exulans*), Southern Fulmars (*Fulmarus glacialoides*), and South Georgia Shags (*Leucocarbo georgianus*) have records of mortality above expected with positive cases being confirmed (Barnyard et al., 2024). More recently unusual South Polar Skuas (*Stercorarius maccormicki*) mortalities were detected from James Ross Island, Antarctica, testing positive for HPAIV H5N1 clade 2.3.4.4. (Bennet-Laso et al., 2024).

Seabirds’ demography is highly sensitive to adult mortality (Lewison et al., 2012; Furness et al., 2000), and some of the affected species already are under extinction risks (e.g. Humboldt penguin, IUCN 2024). The impact of such mortality events on populations of seabirds raised concerns among scientists and conservationists (FAO 2023, Birdlife International 2024). The massive outbreaks scenarios in South America, promoted the early alerts activation in order to guarantee the traceability of the introduction and to determine the spread patterns of HPAIV in Antarctica. Research agencies promptly activated surveillance activities to protect the Antarctic ecosystems, characterized by their unique biodiversity, high endemism, and, at the same time, fragility and vulnerability (Aguado et al., 2024; Bennet-Laso et al., 2024; Rogers, 2007). Surveillance activities have been focused during the austral summer, when the gregarious reproductive behavior of seabirds leads to increased breeding colony densities, which heighten the risk of rapid HPAIV spread, potentially resulting in severe consequences for the Antarctic seabirds.

Evidence of HPAIV presence and susceptibility in Antarctica (Baynard et al., 2024; León et al., 2024; Aguado et al., 2024; Bennet-Laso et al., 2024) has raised significant concerns due to emerging outbreaks and mass die-offs in Antarctic seabird nesting areas. The potential emergence of more virulent viral strains is particularly worrisome, as it could further impact breeding colonies, exacerbating population declines (Tiwari et al., 2024). This threat is especially critical in key seabird conservation areas, such as Antarctica’s Important Bird Areas (IBAs) (BirdLife International, 2015), which support a substantial proportion of global seabird populations or provide habitat for one or more threatened species (Harris et al., 2015).

Among the most vulnerable species are *Pygoscelis* penguins, particularly chinstrap (*P. antarcticus*) and Adélie (*P. adeliae*), which breed in densely packed colonies that often range from tens to hundreds of thousands of individuals (Ainley et al. 1995). Mortality rates observed in other colonial species (10%–60%) suggest a potentially devastating impact on these populations (FAO 2023).

In line with these concerns, our objective was to survey the introduction, spread and ecological impact of HPAIV along the western coast of the Antarctic Peninsula, particularly in areas that are crucial for seabird nesting. This monitoring effort aimed to assess colonies for HPAIV outbreak and evaluate the potential impacts on these breeding sites; the surveillance included both collection of biological samples and behavioral observations at seabird breeding colonies during the 2024/25 season.

## Methods

### Ecological surveillance strategy

Three independent monitoring efforts were conducted during the 2024-2025 summer season: a stationary survey at Harmony Point, Nelson Island, from 23 November, 2024, to 11 January, 2025, and two itinerant surveys; one aboard the Betanzos vessel from January 3 to 16, 2025 and another aboard the Karpuj vessel from January 14 to 15, 2025.

A total of sixteen geographic areas were surveyed along the West Antarctic Peninsula (WAP). These included Antarctic Specially Protected Areas (ASPA) such as Biscoe Point (ASPA No. 139) Harmony Point (ASPA No. 133), Litchfield Island (IBA criteria A4i; ASPA No. 115), and Avian Island (IBA criteria A1, A4i, A4ii, A4iii; ASPA No. 117), as well as other key locations between King George Island and the WAP (Table 1. Fig. 1).

**Table 1:**
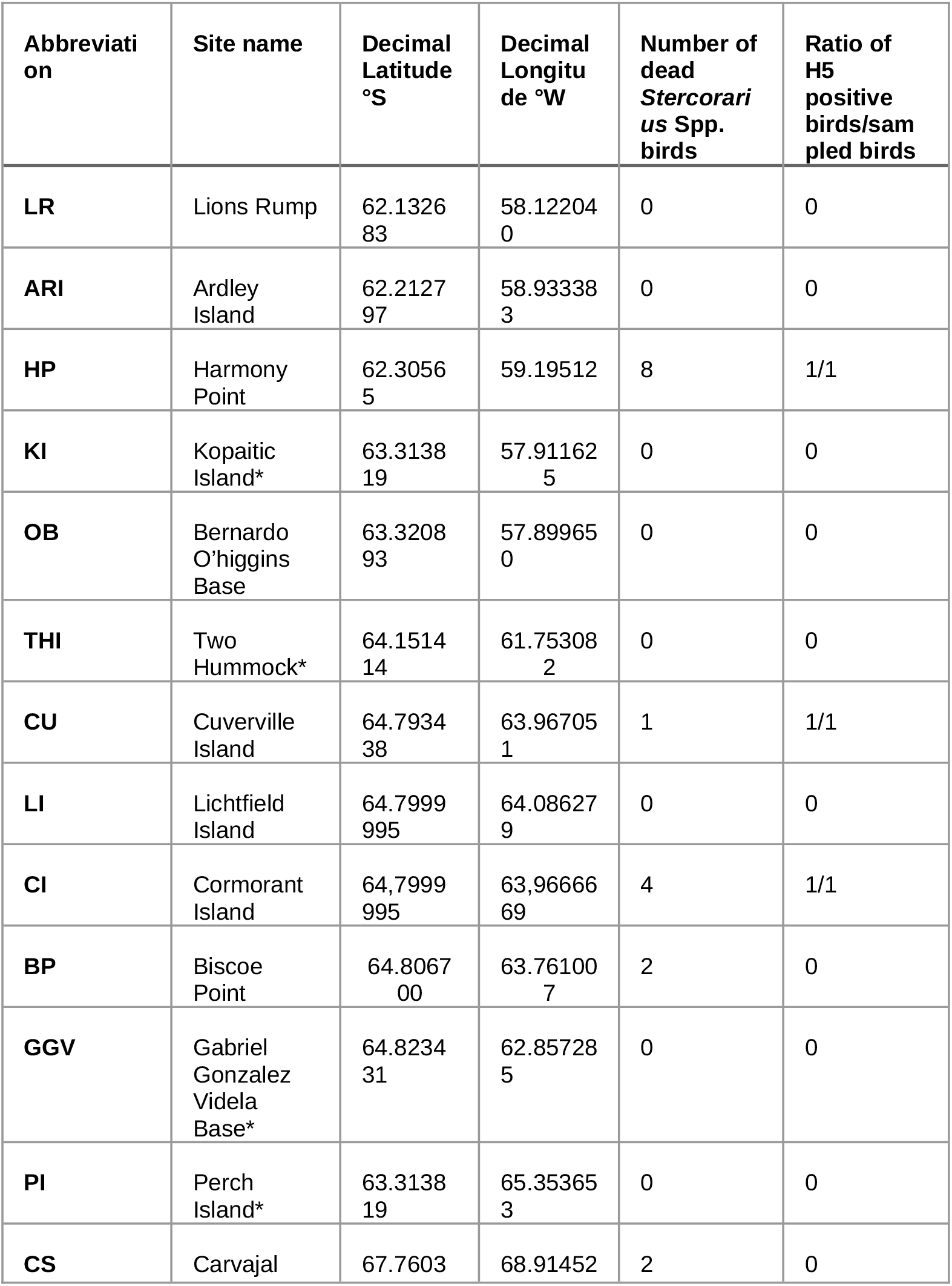
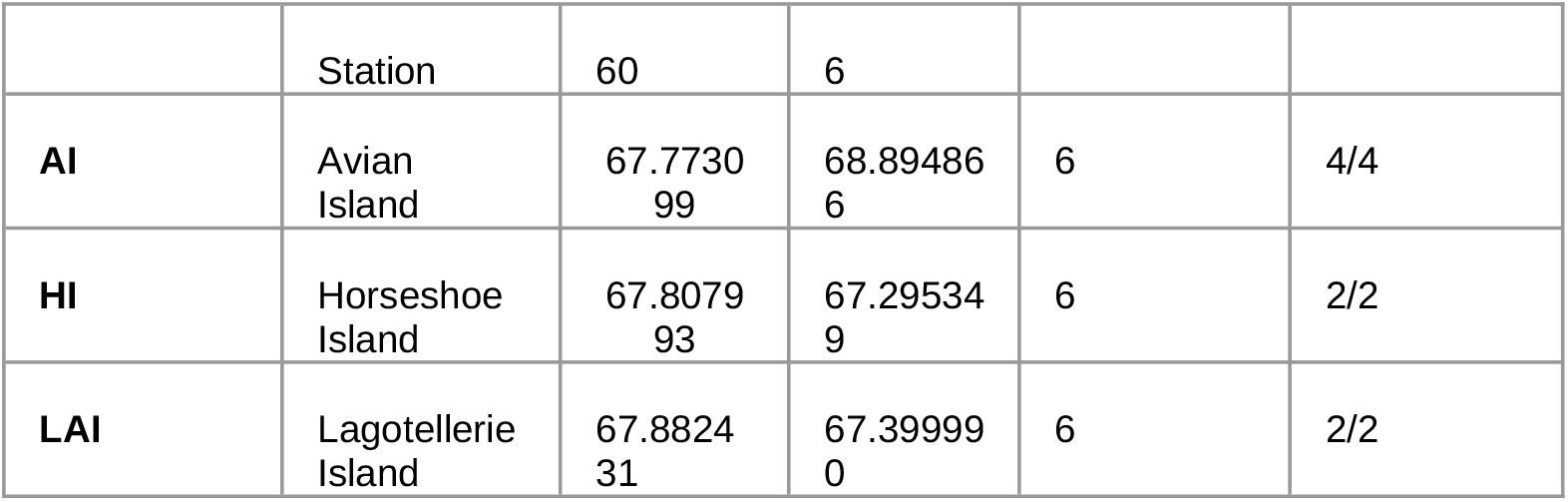
Sampling sites, number of dead and H5 positive adult skua carcasses. Note that not all reported carcasses were sampled. *The location is not reported as a breeding site of skuas.

**Figure 1.**
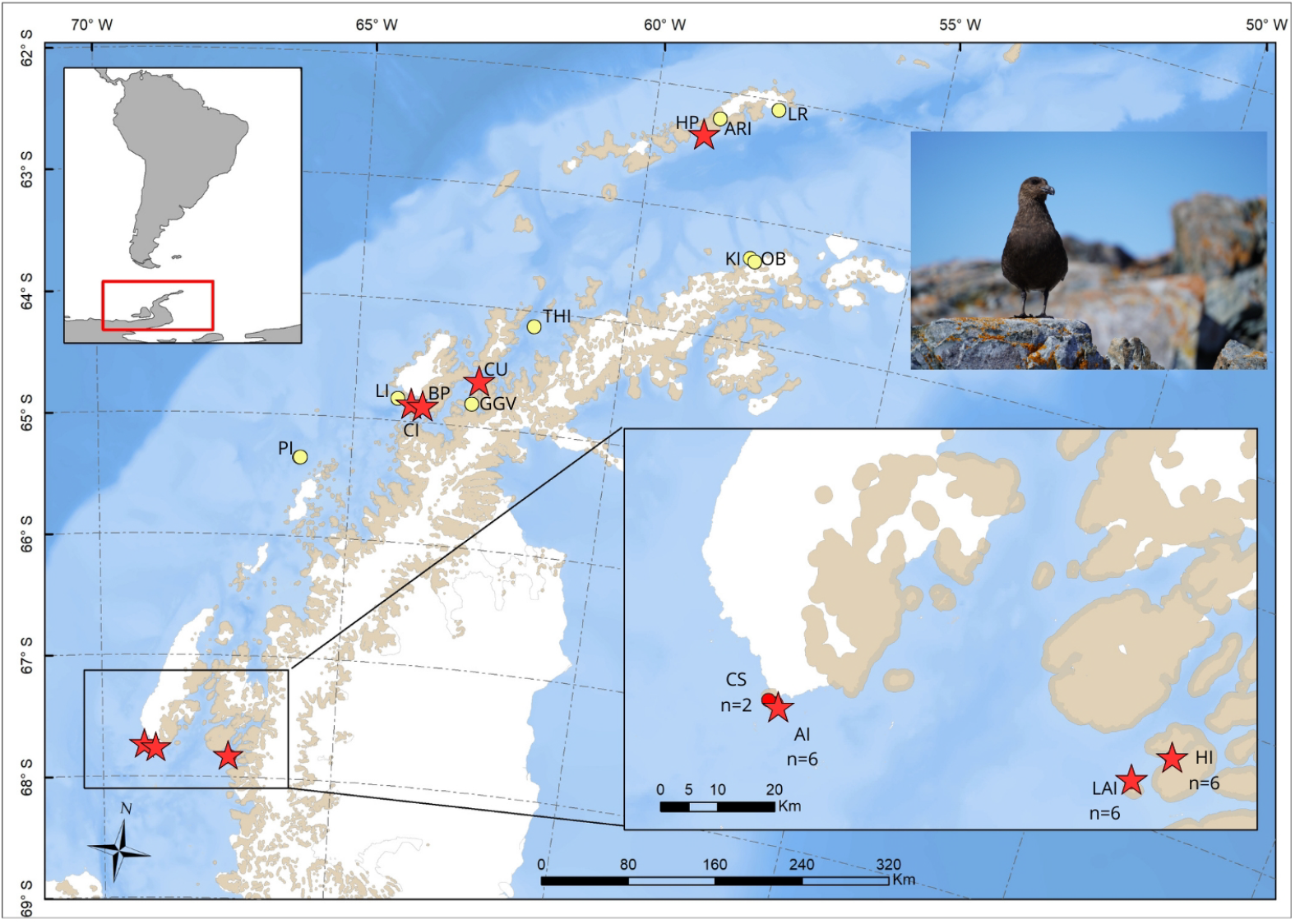
Sampling sites. Unusual mortalities and positive HPAIV cases are denoted with red circles, while no mortalities or suspected cases are indicated with yellow circles. Red stars indicate mortalities and positive cases of HPAIV for new records within the Antarctic Polar Circle. Two of the localities visited, Avian Island (AI) and Lagotellerie (LAI) and Cormorant Island (CI), have ASPA or ASMA categories and all of them are Important Birds Areas (IBA). For abbreviations see Table 1.Photo: [Contanza Barrientos].

From a previous genomic study, all skuas from WAP were previously identified as admixed individuals from hybridization that follows two patterns: i) higher percentage of South Polar skuas (70-96%), and ii) higher percentage of Brown Skuas (56-75%) (Jorquera et al., In press). Although all individuals are admixed in WAP, here we are going to refer to them as South Polar Skuas, or Brown Skuas according to the higher percentage of one of the two species identified in the geographic location.

Visual inspection of birds and colonies was performed to assess the presence of nesting individuals, mass mortality events, or signs of illness suggestive of HPAIV.

For dead skuas, orotracheal, cloacal, and brain swabs were collected for HPAIV detection. Swabs were stored in microtubes, preserved in the Copan eNAT® System, DNA/RNA Shield™ or InhibiSURE™ and then stored in 4°C, -20°C or in liquid nitrogen. In some cases, necropsies were conducted to identify macroscopic lesions consistent with HPAI. Body condition was assessed for all carcasses using a five-point fat scoring system to estimate the subcutaneous fat (Brown, 1996). External palpation and visual inspection were performed to identify traumatic injuries and estimate time of death. If a severely ill and agonizing bird was encountered, it was not handled until it had died.

### Biosecurity measures

All procedures followed strict biosafety protocols adhered to the Antarctic Treaty guidelines and as well as recommendations outlined in the SCAR Biological Risk Assessment for Highly Pathogenic Avian Influenza Virus in the Southern Ocean (Dewar et al., 2023). These protocols were designed to prevent human exposure and minimize pathogen transmission between breeding sites and zoonotic transmission, all samples were transported in inactivation media. All samples collected were obtained under animal capture permits CEC-CAA UC 230103002, CICUA Universidad de Chile: 23722 –VET– UCH and Millennium Institute Biodiversity of Antarctic and Subantarctic Ecosystem 4/CBSCUA/2022. All procedures were approved by Instituto Antártico Chileno (INACH) (permits Nº 00624 / 2024, 00623 / 2024, 00693/2024).

### Molecular detection of HPAIV

Molecular detection of of Influenza A virus was performed on 24 samples derived from 11 skuas at the Laboratory of Molecular Virology, Pontificia Universidad Católica de Chile (LVM-UC). First, samples were processed with TRIzol™ reagent for initial lysis (Invitrogen ™ 15596018) followed by the RNA viral extraction with E.Z.N.A Viral RNA Kit (R6874, Omega Biotek; Pardo-Roa et al., 2023). Then, real-time RT-PCR for influenza a virus was done using the RT-qPCR assay targeting the highly conserved matrix (M) gene of IAV according to WHO (World Health Organization, 2021) recommendations. One positive sample for each bird (CT 21-30) was subsequently tested to determine the Influenza A subtype H5 with the Bio-Speedy(R) Influenza A (H1, 3, 5, 7) RT-qPCR Kit (Bioeksen R&D Technologies, Cat No. FLU-Type) according to the manufacturer’s instructions.

## Results

### Summer 2024-2025 Monitoring Efforts

During the stationary surveillance at Harmony Point, eight Brown Skuas were found dead. Six of them showed evidence of scavenging and lacked sufficient tissue for sampling. During this period, two adult skuas exhibiting neurological signs consistent with HPAI were observed. Both individuals had lethargy and neurological signs which included tremors, opisthotonus and loss of equilibrium (supp video 1). Only one of both birds was confirmed dead, and the samples were collected. That bird remained in a snow melt stream with general and neurological signs which worsened until its death. It took around two days between that bird was first spotted with signs and the confirmation of death. The second bird was found in its nest with neurological signs, loss of equilibrium and maneuvering its wings, and was not found in the following days in its nest.

During the itinerant expeditions aboard the Betanzos and the Karpuj vessel a total of 35 dead skuas were found across the WAP, in 8 sites including Lagotellerie Island (n=6), Harmony Point (n=8), Horseshoe Island (n=6), Avian Island (n=6), Cormorant Island (n=4), Carvajal Station (n=2), Biscoe Point (n=2), and Cuverville (n=1) (Table 1, Fig. 1). No mortality was observed for the other eight sites, however four of them have not been reported as breeding areas for skuas.

Most dead skuas had well-preserved plumage with no visible signs of trauma or injury. The majority were found in a ventral position, with neck postures suggesting possible opisthotonus at the time of death (Figure 2a, b). Two individuals had dried vomit around their beaks (Figure 2c). Carcasses with no signs of predation had an estimated body condition score of 2/5 or 3/5, indicating an acute disease progression. While most dead skuas appear to have died within 48 hours, some carcasses had been scavenged and lacked sufficient tissue for sampling.

**Figure 2.**
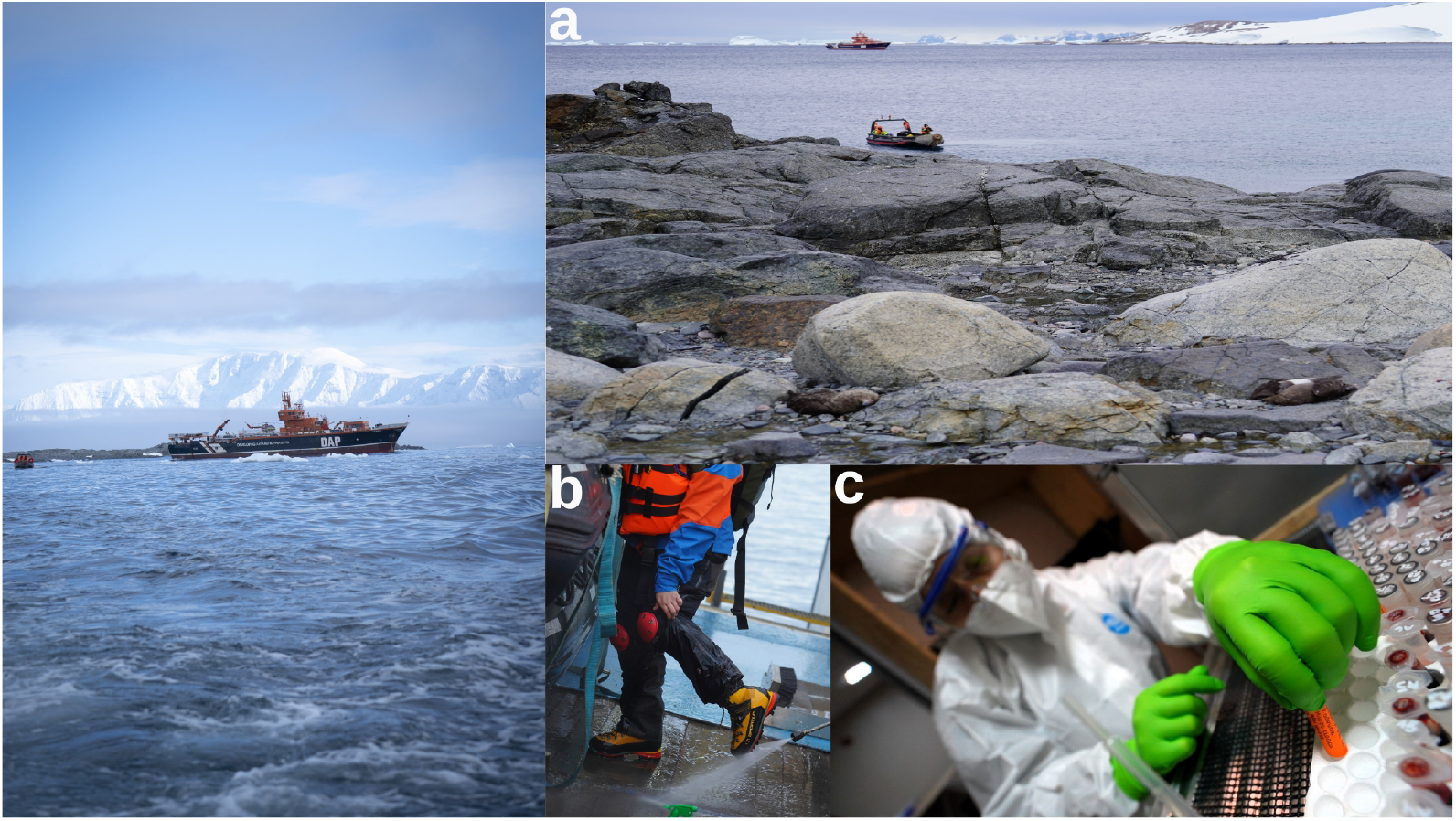
The vessel Betanzos is equipped for conducting scientific expeditions in Antarctica. a) During this expedition, zodiac cruises facilitated the monitoring and observation of seabird colonies. b) The vessel is equipped with biosecurity measures that allow the equipment and clothing to be washed and disinfected. c) A dry laboratory, equipped with the necessary tools and material for stored samples.. Photos: [Constanza Barrientos].

**Figure 3.**
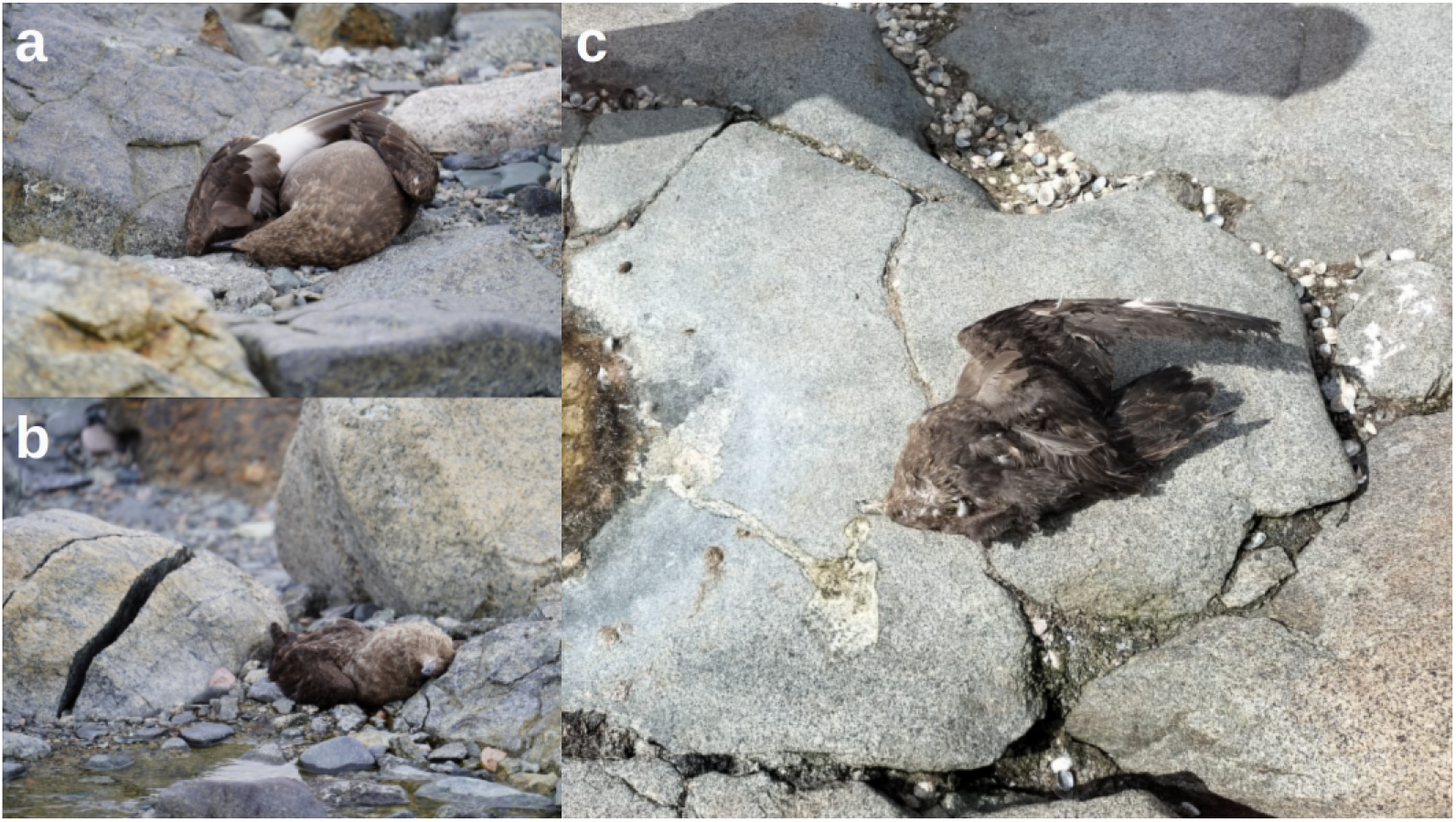
Several dead skuas were observed in pairs near the reported nesting sites in the breeding and rearing areas of the IBA. a-b) Both sampled birds in Horseshoe Island showed neck position indicating a possible opisthotonus sign at the time of death. Birds had no sign of predation and had well-preserved plumage.Photos: [Constanza Barrientos] c) Some animals exhibited dry vomit around the bill. Photo: [Claudia Ulloa-Contreras].

At all sites, typical mortality of *Pygoscelis* spp. and Antarctic Shag (*L. bransfieldensis*) was observed, with some carcasses found isolated along colony edges. In the nesting sites of Southern Giant Petrels, Antarctic Terns, and Snowy Sheathbill, no signs of HPAIV, carcasses, or declines in individuals or nests were detected. These species displayed typical territorial behavior for the breeding season.

### Virus Detection

In total 24 cloacal, orotracheal or brain samples were collected from 11 out of 35 dead skuas. A total of 21 samples were positive for Influenza A gene M, 10 of them were also confirmed as HPAIV A/H5, representing a total of 11 positive birds. The samples of birds detected as positive for HPAIV were collected in Avian Island (n=4), Horseshoe Island (n=2), Lagotellerie Island (n=2), Cormorant Island (n=1), Cuverville (n=1) and Harmony Point (n=1) (Table 1, Fig. 1). Those locations all correspond to admixed South Polar Skuas, except for Harmony Point, which should be considered admixed Brown Skua. All positive samples were conclusive for the identification of A/H5 influenza virus with CT values between 13,976 in brain (Avian Island) to 36,503 (cut off 39). Two samples from Avian Island (SP-3 cloacal and SP-6 brain) and one sample from Cuverville Island (SP-1 brain) exhibited CT values under 19 and only one sample had CT >35 (Lagotellerie Island SP-9 orotracheal). Three samples were detected as undetermined, but at least one swab from the same individual was confirmed. No differences were found on the CT values related to locations or tissue (Table 2).

**Table 2:**
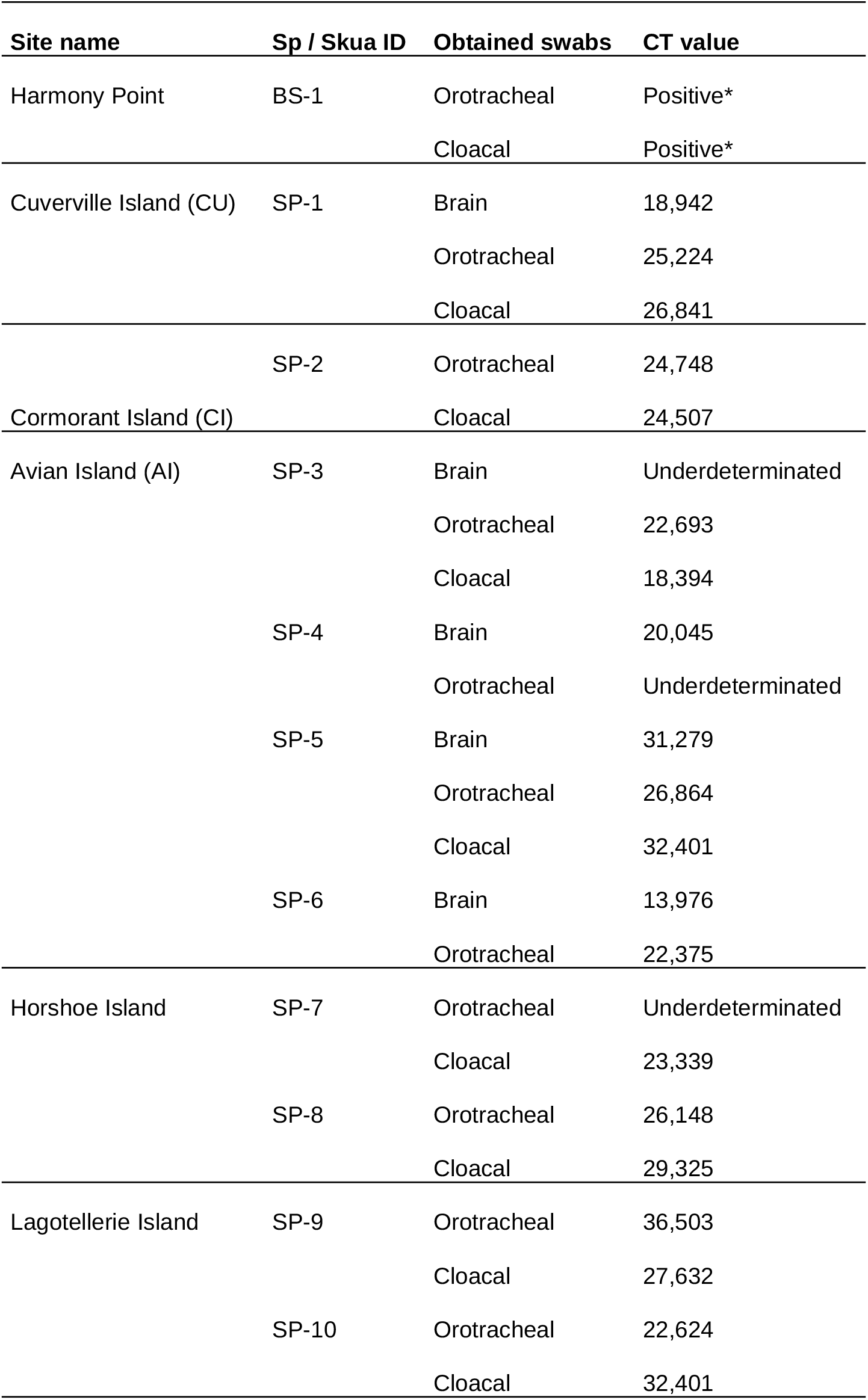
Type of sample obtained from each carcass and its respective Cycle Threshold (CT) value. Sp = Species; Brown Skua (BS) and South Polar Skua (SP). Note that not all carcasses allowed the collection of all three types of swabs. For those carcasses with Positive* result, it was not possible to reobtain CT value.

## Discussion

The detection of the virus in skuas aligns with previous reports of affected species in Antarctica (Bennett-Laso et al., 2024; Aguado et al.,2024;Banyard et al 2024, León et al. 2024). However the detection of HPAIV at its southernmost point in Antarctica is a significant finding that enhances our understanding of the virus’s rapid spread and current distribution on the continent. This study provides the first documented evidence of HPAIV A/H5-associated mortalities and RT-qPCR confirmed cases in the highly migratory South Polar Skua beyond the Polar Antarctic Circle (PAC). The continued spread of the virus beyond the PAC poses a growing threat to Antarctic wildlife and may impact the stability of bird breeding colonies within Important Bird Areas.

In our study, only birds of the genus *Stercorarius* were found dead, exhibiting postmortem signs consistent with HPAIV infection and testing positive to H5, even in areas where breeding colonies of other species were present. The RT-qPCR Cycle Threshold (CT) value is inversely related to viral load, with lower CT values indicating higher viral levels. During season 2024 Bennet-Laso et al. (2024) reports CT values around 19 on brain samples indicating a high viral load on the skua’s population. Here we found a broad range of CT values across samples and individuals, ranging from 14 to 31, with a mean of 24.9. This suggests that viral loads remain important, highlighting the need for continuous surveillance to determine and mitigate the potential ecological outcomes. Both *S. maccormicki* and *S. antarcticus* migrate to lower latitudes during the austral winter. However, *S. maccormicki* undertakes a much more extensive migration, reaching the North Atlantic and Pacific oceans to overwinter (Kopp et al., 2011), whereas *S. antarcticus* typically remains within the southern hemisphere (Krietsch et al., 2017; Delord et al., 2017). This difference in migratory patterns is highly relevant to the spread of HPAIV from the Northern Hemisphere, as *S. maccormicki*’s trans-equatorial migrations expose it to infected regions in the north, increasing the likelihood of the virus being transported back to Antarctic ecosystems. In contrast, while *S. antarcticus* has more localized movements, its interactions with other migratory species could facilitate regional transmission within the Southern Hemisphere. As predators, scavengers and kleptoparasites with high migratory capacity, both species may play a role in facilitating and contributing to the spread of the virus (Gorta et al.,2024)

Moreover, South Polar and Brown Skua overlap their distribution across 500 km of the Antarctic Peninsula (Ritz et al., 2006,2008). Comparative genomic study indicates significant genetic admixture along the Antarctic Peninsula between Brown, Chilean, and South Polar skuas, with a low population structure within breeding colonies in WAP (Jorquera et al., 2025). These high levels of gene flow may be further contributing to the virus transmission across populations and species, both within and between sub-Antarctic and Antarctic colonies.

Our observations during the 2024-2025 Antarctic expeditions detected a higher number of dead skuas within the Antarctic Circle, where the introgression pattern between South Polar Skua and Brown Skua changed, decreasing significantly the percentage of admixture with Brown Skua. Interspecific gene flow could serve as a significant source of novel, adaptive variation (Hessenauer et al., 2020). Specifically, multiallelic balancing selection, particularly through rare allele advantage, may facilitate adaptive introgression by conferring immediate selective benefits, aiding the establishment of introgressed alleles in recipient populations (Hedrick et al., 2013; Fijarczyk et al., 2018).

For populations unable to generate an effective response against HPAIV H5N1, the increasing threat of EIDs raises concerns about population bottlenecks that could lead to local extinctions. This risk is particularly critical for endemic reproductive species in valuable regions such as South Polar Skua in Antarctica. In this unique environment, population bottlenecks, along with declines and recolonization events, can disrupt population structure, leading to fragmentation and reducing effective population sizes, ultimately accelerating the loss of genetic diversity across multiple populations.

Population changes in both species do not follow a similar demographic trend. At Ryder Bay, one of the southernmost distributions on the Antarctic Peninsula, a long-term population density study shows an overall increase of breeding pairs and occupied territories of *S. maccormicki* (Phillips et al., 2019). Further north, at Harmony Point, the most recent study indicates that population sizes of both species have remained relatively stable over the last 25 years (Santa Cruz & Krüger, 2023). In the more northerly regions, population trends appear to differ. At Signy Island in the South Orkney Islands, the breeding population of *S. antarcticus* has increased, whereas *S. maccormicki* has declined from 10 breeding pairs to just one (Carneiro et al., 2016), and in Elephant Island there are records of recent decreases in at least one breeding site (Stinker Point, Petry et al. 2018). These variations may result from multiple factors, including dietary differences and the competitive abilities of each species in each area (Carneiro et al., 2016; Reinhardt et al., 2000; Ritz et al.,2006). Despite these regional differences through the latitudinal distribution, population growth in the coming years seems to be unlikely, as birds might be occupying all the available breeding niche on a given site (Santa Cruz & Krüger, 2023).

According to the International Union for Conservation of Nature (IUCN) both species are categorized as “Least Concern”. However, while South Polar Skua has a stable global population trend, Brown Skua is decreasing even before the emergence of the HPAIV (BirdLife International, 2018). Future scenarios for these species suggest that Brown Skua might experience a contraction of its northern distribution and an expansion of the southern distribution which would increase the hybridization area with South Polar Skua (Jorquera et al., in press). In this context, both species can be strongly impacted by HPAIV outbreaks and consequent mortalities. On one hand, Brown Skua could experience an accelerated population decline, while South Polar Skua may undergo a reduction in population density due to increased hybridization and mortality. The threat could be even more significant, considering that many local adaptations involve immune system genes (Jorquera et al., in press).

For the breeding population of Brown Skua at Harmony Point, Santa Cruz and Krüger (2023) estimated approximately 70 to 80 breeding pairs, with no evidence of decline in the last season 2023-2024. However, during the 2024-2025 season, the report of eight Brown Skuas found dead at this site represents an important number and could have a significant impact on the colony. While no decline in the number of breeding pairs was observed, changes in the breeding individuals within the area cannot be ruled out if new recruits replaced those that perished.The number of recorded skua deaths this season exceeded the usual 1 to 3 adult deaths typically reported in the area. However, this is likely an underestimate, as monitoring was conducted over a limited period and did not account for at-sea mortality. Notably, as in previous studies, mortality tends to be underestimated when relying solely on carcass searches (Ward et al., 2006)

Infectious diseases have been emerging at unusually high rates in recent years (Daszak et al., 1999; Epstein 2001) and have been linked to species extinctions (Daszak et al., 2000; Harvell et al., 2002). Epizootic spillover outbreaks pose not only a significant risk to wildlife but also a threat to human health due to the potential for pathogens to spill back (Daszak et al., 2000). The disruption of proper ecosystem functioning and the essential services it provides for human well-being are among the key impacts of biodiversity loss (Mace et al. 2012), that eventually could lead to species extinctions. Mahon et al., (2024) found that changes in biodiversity patterns could drive the rise of EIDs even more than anthropogenic factors like pollution and climate change. The complex relationship between how EID dynamics influence changes in host biodiversity and vice versa complicates the approach to addressing this issue. Its consequences rank among the top five factors that can drive extinction risk in the United States (Wilcove et al., 1998). According to Keesing et al (2010) while higher biodiversity can reduce disease risk through the “dilution effect,” it can also increase it via the “amplification effect.” The complex interplay between community composition, ecosystem diversity, and EIDs forms a dynamic system that remains poorly understood (Cunningham et al., 2017). Our results alert about demographic impact on a key apex predator in the Antarctic ecosystem, which could, in turn, alter the demographics of other species within the same trophic chain. During the breeding season, Brown Skua primarily prey on penguin chicks and eggs, whereas South Polar Skua has a diet with higher consumption of fish and krill (Reinhardt & Hahn, 2000). These differences might also affect the eco-epidemiology of the HPAIV on the WAP, through differences in the predator-prey and multiple host species dynamics (Roberts & Heesterbeek, 2018; Sander et al., 2007). The Antarctic continent has been facing a reduction of seabird population such as Chinstrap Penguins, Adelie Penguins and Cape Petrel (*Daption capense*) (Carlini et al., 2009; Braun et al., 2021; Krüger, 2023; Sander et al., 2007, Talis et al 2023, Petry et al. 2018). The impact on community diversity composition through multifactorial effects could lead to an increase in the transmission and incidence of infectious diseases (Keesing et al., 2010).

It is noteworthy that, although being a highly migratory species and a potential candidate for transpolar virus transmission, the Antarctic Tern does not appear to be affected by HPAIV, at least not by notable mortalities, even though it nests near areas where outbreaks and mortalities occurred. Similarly, the Giant Petrel, an Antarctic scavenger with a more limited migration range, has shown possible signs of HPAIV in sub-Antarctic environments. However, no positive samples have been obtained from this species, and no unusual mortalities have been observed in its populations (Banyard et al., 2024). Likewise, the Snowy Sheathbill, another scavenger and kleptoparasite previously reported as positive for HPAIV in Antarctica (Aguado et al., 2024), showed no signs of unusual mortality or HPAIV infection. These observations pinpoint the varying impacts of the virus across different seabird species

Given this unexpected mortality in Specially Protected Areas beyond 66.5°S and the rapid spread of the virus, it is crucial for IAATO and SCAR authorities, scientific researchers, conservation managers, and international organizations to implement urgent measures to improve the surveillance of the HPAIV in order to minimize the adverse effects of the outbreak. Important Bird Areas (IBAs) in Antarctica, such as those near Margarita Bay, play a critical role in global seabird conservation efforts. The confirmation of HPAIV in these IBAs underscores their vulnerability and highlights the need for targeted conservation measures. A lack of preventive action in Antarctica could cause irreversible damage to its biodiversity, making proper management crucial now to protect the ecosystem and species (Handley et al., 2021). Strengthened biosecurity measures, even in the absence of direct scientific research on birds, will be critical in managing the HPAIV threat in IBAs within fragile ecosystems such as Antarctica. Enhancing monitoring efforts, including colony densities counts will provide a better understanding of the outbreak status and its impact on the Antarctic seabirds. Long term systematic monitoring of biodiversity in Antarctica has been suggested to be crucial for conservation practice (Pertierra et al., in 2025). Continuous monitoring of migratory birds through movement ecology analysis will allow for greater insight into the sites of infection and help identify surrounding variants in the Antarctic region, as well as the virus’s evolutionary potential and its relation to pathogenicity.

## Acknowledgments

This study was funded by ICM-ANID ICN2021_002 Millennium Institute BASE, the Oceanographic Institute, Foundation Albert I, ICN2021_044 – CGR, INACH RT-30-22 and National Institute of Allergy and Infectious Diseases, National Institutes of Health, Department of Health and Human Services, Centers of Excellence for Influenza Research and Response (CEIRR) under Contract No. 75N93021C00017 Option 18A, Brazilian National Council for Scientific and Technological Development (CNPq), under the Project number 440901/2023-5, and Instituto Antártico Chileno (INACH) Programa Areas Marinas Protegidas (AMP 24 03 052). We are very grateful to the all team of ICM-ANID ICN2021_002 Millennium Institute BASE. We deeply thank the Betanzo Crew, the Capitán José Reyes, Edgardo Barrios Villouta; the INACH logistic staff, in particular Alejandro Font and Pablo Espinoza; Carolina Márquez and Constanza Barrientos for their help on the fieldwork and Melisa Gañan Mora for the map.

